# Elucidating the complex organization of neural micro-domains in the locust *Schistocerca gregaria* using dMRI

**DOI:** 10.1101/2020.01.17.910851

**Authors:** S.S. Shahid, C.M. Kerskens, M. Burrows, A.G. Witney

## Abstract

To understand brain function it is necessary to characterize both the underlying structural connectivity between neurons and the physiological integrity of these connections. Previous research exploring insect brain connectivity has used microscopy techniques, but this methodology is time consuming and cannot be applied to living animals and so cannot be used to understand dynamic physiological processes. The relatively large brain of the desert locust, *Schistercera gregaria* (Forksȧl) is ideal for exploring a novel methodology; diffusion magnetic resonance imaging (dMRI) for the characterization of neuronal connectivity in an insect brain. The diffusion-weighted imaging (DWI) data were acquired on a preclinical system using a customised multi-shell diffusion MRI scheme. Endogenous imaging contrasts from the averaged DWIs and Diffusion Kurtosis Imaging (DKI) scheme were applied to classify various anatomical features and diffusion patterns in neuropils, respectively. The application of micro-MRI and dMRI modelling to the locust brain provides a novel means of identifying anatomical regions and connectivity in an insect brain. Furthermore, quantitative imaging indices derived from the kurtosis model that include fractional anisotropy (FA), mean diffusivity (MD) and kurtosis anisotropy (KA) could, in future, be used to quantify longitudinal structural changes in neuronal connectivity due to environmental stressors or ageing.

## Introduction

A key challenge of neuroscience is understanding the emergence of behaviour from neuronal activity. The idea of neuronal circuit formation sharing common design principles across different species to generate behavioural equivalent outputs has had a long history since the pioneering work of Ramon y Cajal and his early observations of the similarity in organization between insect and human visual processing ^1^. Interestingly, regardless of the number of neurons that comprise the brain of an animal and morphological differences, evidence suggests that the basic principles underlying neuronal connectivity are similar ^2,3^.

The emergent field of connectomics develops the premise that the understanding of how complex brain function must be related to its structural underpinnings ^4^. Initial studies of neuronal connectivity were in *Caenorhabditis elegans*, an animal with only 302 neurons with simple behaviours^5^. However perhaps both the simplicity of the behaviour of *C. elegans* and its decentralised nervous system make it an inadequate model animal to link neuronal circuitry to behavioural output in a way that can be generalizable across species. Recent work in the *Drosophila melanogaster* connectome has greater potential as insects have long provided important model animals for understanding the relation between neural circuitry and behaviour using electrophysiological techniques ^6^. Whilst the insights gained from electrophysiology have been essential in understanding how the processing in discrete neural circuits relates to simple behaviours, these techniques may be limited in the insights they can provide about the neuronal network as a whole. Network level understanding could be critical in understanding the emergence of more complex behaviours that inevitably rely on the integration of multiple discrete neuronal circuits. Linking electrophysiology and connectomics could prove to be a very powerful methodology in understanding complex adaptive behaviour^7,8^.

The behaviour of the fruit fly *Drosophila melanogasta*, is far more diverse than that of *C. elegans* and the animal has a correspondingly more complex nervous system, with an estimate of 135,000 neurons. Shih et al (2015) produced a connectome of the *Drosophila* nervous system, using 12995 images of projection neurons collected in ‘FlyCircuit’^9^. The neurons were found to connect 43 local processing units resulting in the formation of five distinct modules: left visual, right visual, olfactory, mechano-auditory and pre-motor module. Terminology based on the graph theory origins of connectomics have been used to characterize similarities in neuronal network structure across phyla. For instance fly brain connectomes demonstrated the presence of ‘rich club’ organisation in insect brains, that is ‘nodes’ that are extremely linked and can be measured by the overall strength of connections^9^. Evidence of this and ‘small world’ architecture in the formation of neuronal processing networks has been observed in mammalian brains ^10^. This form of analysis can be used to argue that whilst the mammalian brain is far more complex than the insect brain, the key characteristics of its network formation are essentially equivalent, thus providing further support for the concept that insect brains are excellent model systems for understanding the neuronal basis of complex, adaptable behaviour ^4,11^. Whilst these approaches from graph theory provide an overall computational approach to understanding neuronal functioning they cannot be easily related to biological changes that occur in fibre tracts during development, ageing or injury.

Thus far the construction of the complete connectome using *C. elegans* and *Drosophila* has relied on the use of semi-automatic processing to produce reconstructed electron microscopy images^12^. The resolution of these technique enables comprehensive synapse level connectivity to be determined in an adult fly brain ^11,13^. This remains a time realistic approach for these animals, given their simpler nervous system, and one that has begun to characterize relationships between neuronal architecture and behaviour. However, these techniques are limited ^14^. The reconstruction of electron microscopy images remains very time consuming process even in these animals with simpler nervous systems. The process of reconstruction itself is also susceptible to error. Microscopy further necessitates the animal being dead and so presents a static image of neuronal connectivity ^15^. This approach also limits sample size, with the methodology focused on generating a standard connectome for the species rather than providing the possibility of a comparative assessment of differences across animals. To truly understand the link between neuronal networks and adaptable behaviours the techniques for characterizing the network may need to be applied longitudinally over an animal’s lifespan. Further it would be highly advantageous for a visualization technique to be able to quantify micro-structural changes that occur that may reflect damage, ageing or adaptive plasticity in the nervous system. Such a technique would enable mechanistic questions regarding the causes of any alterations in the microstructure of the brain to be addressed.

In parallel to the connectome research in simpler animals, the same interest in examining neuronal connectivity has developed in mammals. In these studies the technique applied has been magnetic resonance imaging and the modelling of the diffusion weighted imaging (DWI) via diffusion tensor imaging (DTI) ^16^. The body of all animals, including insects, has a high water content, and in a biological tissue water molecules are in a constant state of random motion. dMRI utilizes the principle that differences in water diffusion in tissues can be used to indirectly infer underlying anatomy via the direction of preferential diffusion and from this provide unique information on their microstructural architecture ^17^. dMRI provides diffusion estimates for each voxel in a series of DW images. DTI and higher order diffusion schemes can then describe the estimated water self-diffusion in different dimensions by a diffusion tensor or more advanced representation schemes. This tensor mathematically represents how the composite architecture of the body provides structural barriers differentially restricting the directions of water diffusion. The directionality of water diffusion that emerges due to these biological barriers in the body is known as anisotropic or directional diffusion, in contrast to isotropic diffusion where there is no directionality. In the brain the biological barriers to water diffusion include axons, cellular membranes and proteins. Therefore, DTI allows the visualization of fibre pathways and connectivity. Importantly, DTI does not directly visualise axonal connections, but is a statistical inference of structural connectivity ^18 19^. However DTI is limited by assumptions intrinsic to it regarding the Gaussian distribution of water self-diffusion within the body, a simplification that will reduce the accuracy of the detail that the method can provide about the structures. Diffusion Kurtosis Imaging (DKI) adds additional dimensionality to the model of diffusion and attempts to capture the non-Gaussian distribution of diffusion to give better insights into differences in the barrier for molecular diffusion and from this provide a more accurate inference of the structure of the underlying tissue ^20^.

The DTI and DKI methods, as well as inferring the underlying microarchitecture that is creating the barriers to diffusion, enables the quantification of differences between different regions of the brain in terms of how they impact to the diffusion of water. There is evidence that measures derived from DWI via DTI are sensitive enough able to detect pathological changes in axonal architecture that are associated with the osmotic swelling occurring as a consequence of alterations in membrane polarization ^21^. Such axonal changes are thought to be indicative of Wallerian degeneration or axonal self-destruction that occurs in disease and after trauma ^22^. Understanding the mechanisms behind the altered architecture of axons is important as this pathological process is thought to be characteristic of many neurodegenerative diseases. Although there are morphological differences between insects and vertebrates, similarities at the cellular and molecular level, with both neural signaling and the innate immune response highly conserved from insects through to mammals^23^, have led to insects being proposed as important models for understanding the biological pathways underlying axonal self-destruction. Fractional anisotropy (FA) is the most widely applied metric that quantifies differences in water diffusion by biological barriers, and provides a measure of strength of directionality of diffusion in a given region. Differences in FA have been used widely in DTI studies of the human brain and have found to be of predictive value in characterizing the severity of pathological processes in the CNS due to trauma or neurodegenerative diseases ^16,20^.

DTI or DKI has not previously been implemented on insects but the application of dMRI and DWI scans provides the advantage of providing a much faster means of visualising the locust brain microcircuitry and neural tracts. To be able to use DTI or higher-order diffusion schemes ^24,25^ to characterize the neural connectivity of simpler species up to more complex animals will allow us to build upon knowledge in the growing field of comparative connectomics ^26^. A recent study applied dMRI to the squid brain, both confirming the presence of connectivity previously established by microscopy techniques, and also identifying many previously undescribed pathways^27^. Further the application of dMRI enables the novel opportunity for quantification of indices that may reflect structural alterations that are occurring due to disease, injury or plasticity. Due to the known commonalities at the cellular and molecular level between insects and vertebrates, the use of these quantitative measures in an insect model in combination with genetic tools ^28^ has potential in the future to provide insights into axonal biology *in vivo*.

## Materials and Methods

### Sample preparation

Three female 5^th^ instar desert locusts, *Schistercera gregaria* (Forksȧl) (Blades Biological, Kent, UK) was decapitated by cutting rostral to the pronotum. The locust head was then put in standard locust physiological saline ^29^ before stabilizing in a 10 mm falcon tube, with the head immobilized by plastic holders. The antennae were removed from the locust head to facilitate good stabilization of the head within the falcon tube.

### MR imaging

The DWI data were acquired at the bore temperature on a horizontal bore 7T Biospec micro-MRI system (Bruker, Etlingen Germany) equipped with shielded gradients (maximum gradient strength = 770 mT/m, rise time = 115 μs) and ^1^H mouse cryogenic surface coil (cryoProbe, Bruker Biospin). The acquisition was conducted in coronal plane with the following acquisition parameters: 2D spin echo based diffusion sequence with unipolar diffusion sensitizing pulse field gradients placed symmetrically around the 180° RF pulse. TE/TR = 17.628 /1000 ms; FOV = 7.5×10 mm; matrix size 96×128; slice thickness 0.781 mm; voxel size = 78.125 μm^3^. No inter slice spacing; number of slices = 35; no fat suppression; δ/Δ = 3.1/8.502 ms. The prescribed b-value = 800, 1800 and 2500 s/mm^2^, 6 directions for the first shell (b=800 s/mm^2^), 8 directions for the second shell (b=1800 s/mm^2^) and 12 directions for the third shell (b=2500 s/mm^2^). In total, there were 1 b0 image per shell and a total of 26 diffusion sensitizing gradient directions. The entire DWI volume was collected in a total scan time of ≈ 24 hours.

### Diffusion MRI Modelling

In a pulsed gradient spin-echo (PGSE) diffusion sequence ^30^, if the diffusion-encoding gradient with amplitude *G* has infinitesimally short pulse duration *δ* compared to the diffusion time *Δ*, then the displacement of water molecules during this short diffusion gradient time can be ignored. Therefore, under the assumption of narrow pulse approximation (δ << Δ) ^31^, the signal attenuation due to the diffusion-encoding gradient is given by:

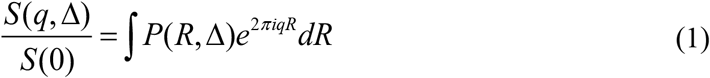

Where *q*=*γGδ/2π* is a wave vector ^32^, *γ* is the gyromagnetic ratio, *R* is the dynamic displacement (displacement of spins during the allowed diffusion time *Δ*), and *P(R)* is the ensemble average propagator (EAP) and it provides averaged estimate of the diffusion environment. The diffusion profile of a complex structure can be obtained using the inverse Fourier transform of the signal with respect to the wave vector ^32^ and the excess kurtosis can be calculated using the following relation ^33^:

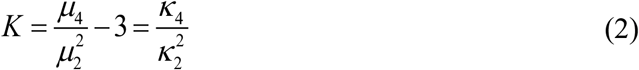

Where *μ_i_*= ∫ *R*^*n*^*P*(*R*)*dR*, the cumulants (*κ_i_*) can be described in terms of moments of probability distribution (*μ_i_*). The first three cumulants (*κ_1−3_*) are equal to the first three moments (*μ_1_*_−3_). *μ, μ_2_* and *μ_3_* are the mean, variance and the skewness of the distribution, respectively. The fourth cumulant is related to the kurtosis as shown in eq. 2. Diffusion Kurtosis can be derived from dMR signal by using the Fourier relationship between the attenuated signal and the EAP. The logarithm of the diffusion signal can be expanded as a summation of the cumulants *κ_n_* of *P(R)^33^*:

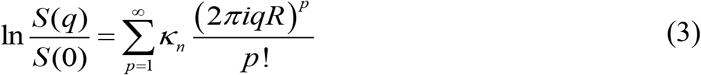

Under the assumption that diffusion is symmetrical (symmetric EAP), the phase of the attenuated signal can be considered zero, therefore, all odd order cumulants are null, i.e.,

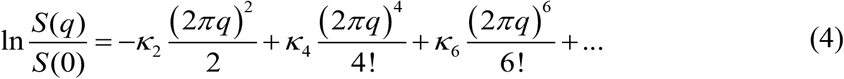

For isotropic Gaussian diffusion in time *Δ*, the diffusion coefficient can be expressed as: *D*=*κ_2_/(2Δ)* ^24^, and by substituting it in eq. 2, the fourth cumulant can be written as: *κ_4_*=*4KD^2^Δ^2^*.

Under the assumption of PGSE, the diffusion weighting parameter ‘b-value’ is defined as:

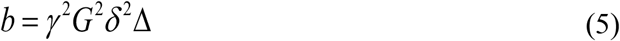

Using the relations of the second and fourth cumulants and eq.5, the signal attenuation can be approximated by the quadratic exponential kurtosis model after truncating eq. 4 to the second term:

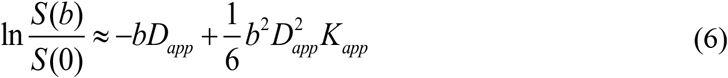

Where, *b*≈*Δ(2πq)^2^*, and when *δ* ≈ *Δ* (violation of narrow pulse approximation), the effective diffusion time ‘*τ*’ (*τ* = *Δ – δ/3*) should be used in eq. 5 instead of *Δ*, i.e.,

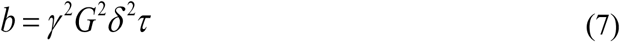

By measuring the attenuated signal with multiple b-values, it is possible to estimate the apparent diffusivity (*D_app_*) and apparent diffusion kurtosis (*K_app_*) along a specific diffusion direction by fitting eq.6 ^34^. The PDF of the diffusion kurtosis model (eq. 6) is a Gaussian distribution with mean (*D_app_*) and variance 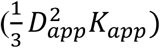. Therefore, kurtosis can be used to estimate the degree of heterogeneity of the underlying structural environment.

For b-values that are significantly small (depending upon the type of tissue) and if a voxel contains a single compartment exhibiting homogeneous T2 relaxation, then for unrestricted Gaussian diffusion, the higher terms of eq. 4 can be neglected and eq.4 takes the form of a well-known mono-exponential (DTI) model:

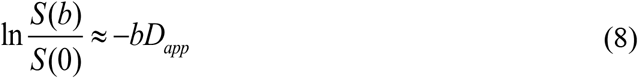

By fitting eq.8 over a range of b-values, *D_app_* can be estimated. However, if the range of b-values is very small (maximum b-value is too low) then the attenuation in the signal intensity will be very low and the estimation of *D_app_* will be prone to noise. On the other hand, if the range is very large (max b-value is very high), then the estimation of diffusivity will incur a systematic error due to the omission of higher terms from eq. 4.

### Diffusion MRI data processing

The noise profile of DWI at higher b-values is non-Gaussian, therefore, to reduce the influence of noise, NLMeans filter with rician noise estimation was used for each dwi dataset ^35^. The noise compensated datasets were then processed to correct for Gibbs ringing artefacts ^36^ and subsequently corrected for eddy current induced distortions ^37^. Since, mono-exponential derived diffusion indices are highly susceptible to diffusion gradient strength and the b-value, therefore, in this study, a Kurtosis model (eq. 6) was used to derive all diffusion-based scalar indices ^38,39^. DESIGNER which is a python/matlab based API was used for preprocessing and to derive DTI and DKI derived imaging indices (https://github.com/NYU-DiffusionMRI/DESIGNER). Imaging indices such as fractional anisotropy (FA), mean diffusivity (MD), mean kurtosis (MK) and kurtosis anisotropy (KA) were derived from the kurtosis tensor ^24,25^. FA measures the degree of diffusional anisotropy and has a range of 0 and 1 as it derives from a ratio of random to highly-directional diffusion from the diffusion tensor. Higher FA values suggest higher directionality of diffusion. MD represents the diffusion rate within a voxel, with higher values representing increased diffusivity or isotropy; MK is the average diffusion kurtosis along all directions, with higher values representing increased restriction to molecular diffusion (non-Gaussian) along these dimensions; KA estimates the variability in the kurtosis and is derived from the standard deviation of the kurtosis. KA has a range from 0 to ∞ and it captures diffusional anisotropy with the additional dimensionality that is provided by the kurtosis model.

For tractography based assessment, DSIStudio ^40^ was used with the following parameters: FA tracking threshold 0.15, fibre length range [0.18 50] mm, angle threshold =45 degree, step size = 0.05mm, tracking algorithm =Euler and interpolation method = cubic. For each dataset 5000 fibre tracts were generated and exported to TrackVis ^41^ compatible format for visualization.

## Results

dMRI enabled anatomical regions to be identified from the DWIs and subsequently tract connectivity in the locust brain to be extracted based on the diffusion direction over a series of DWIs. Figure 1 shows multiple anatomical features of the inset in coronal, axial and sagittal planes. For each plane, the top row shows the anatomical features using T2-W contrast, whereas the bottom row shows the anatomy using the contrast generated from the averaged DWIs.

**Figure 1:**
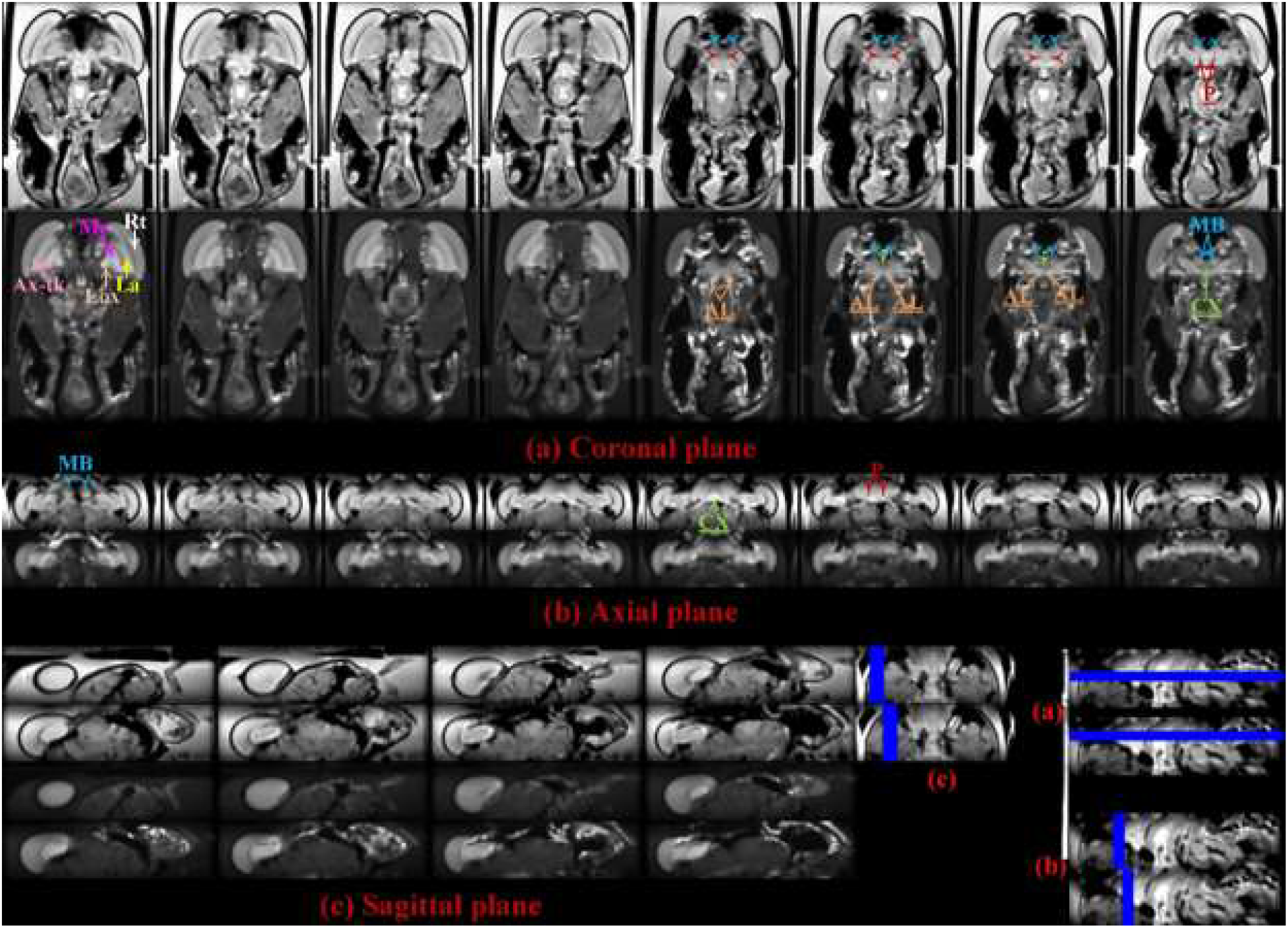
Anatomical images (T2-W and averaged DWIs) to show various regions in the head. (a) Coronal plane (top row T2-W, bottom row averaged dwi). Rt: retina; La: lamina; Ax-tk: axonal tracts; MB: mushroom body; Me: medulla; AL: antennal lobe; P: peduncles; Lox: lobula complex. (b) Axial plane (top row T2-W, bottom row averaged dwi). (c) Sagittal plane (top row T2-W, bottom row averaged dwi).

In addition to the anatomical features, quantitative metrics can be extracted from the DWIs, including FA. The use of these metrics have the potential for the visualization of the locust brain to be quantified and therefore used to correlate with observed physiological changes. Figure 2 shows a coronal slice with a number of contrasts derived from the acquired DWIs and the Kurtosis model. In b0 or T2-W image (Figure 2-a), the highest contrast is from fat bodies, whereas, in Figure 2 (d) the signal from fat bodies is suppressed and regions of hindered/restricted diffusion displayed high contrast ^42^. Using this multi-contrast approach, it is possible to identify various regions, such as Retina (Rt), Lamina (La), Medulla (Me), peduncles (P), axonal tracts (ax-tk), Mushroom body (MB), Lobula complex (Lox) and antennal lobe (AL) ^43^. Figure 2 (b) shows the FA modulated directionally encoded color (DEC) map. Using DEC maps, it is possible to identify several synaptic microdomains. Axonal tracts, which were hypo-intense in both T2-W and averaged DWIs, show blue contrast indicating the out of plane fibre orientation and axonal tracts in the MB indicating vertical orientation (Figure 2-b). The DEC map can also identify a number of regions, such as AL, Lox and the synaptic centers of the optic lobe. Figure 2 (c and e) therefore provide a more quantitative insight of the microstructural environment. The FA map shows high contrast in Rt, La, Me and Lox. This would be indicative of these regions having a high directionality which would be consistent with the visual processing pathway, from the retina through the optic lobes towards the central brain. In contrast, the optic lobe regions are hypo-intense in mean diffusivity map, which is thought to relate to higher levels of structural complexity. This finding would reflect the layering of synaptic networks, with each neuropil known to consist of both columnar arrangement of neurons and also local interneurons^44^.

**Figure 2:**
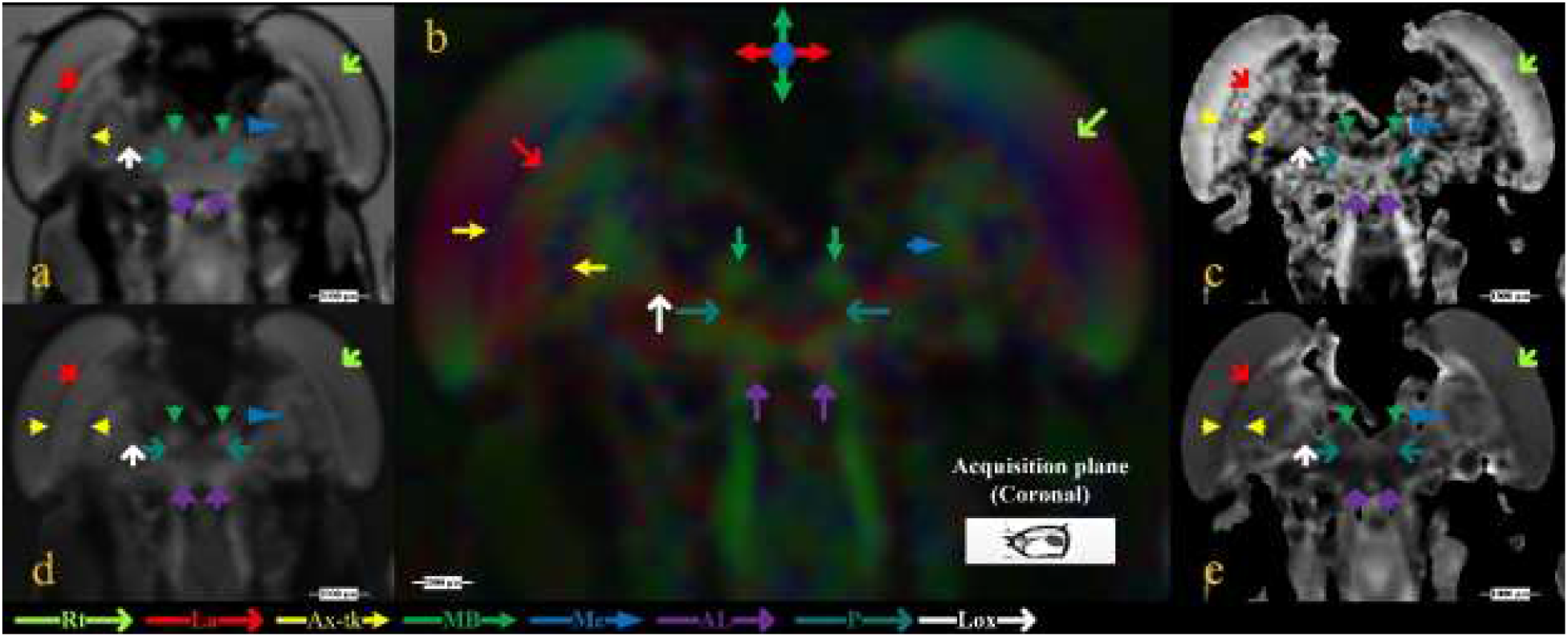
Coronal slice, anterior view from sample no.2 illustrating various anatomical structures and their respective diffusion profiles. (a) b0 or T2-W image; regions with longer transverse relaxation (T2-relaxation) are hyper-intense, whereas, regions exhibiting shorter transverse relaxation are hypo-intense, for example, cornea has the shortest T2-relaxation and La has comparatively longer T2-relaxation time, (b) Fractional anisotropy (FA) modulated directionally encoded color (DEC) map, which can be used as a visual representation of the orientation of the primary eigenvector. (c) FA map highlighting regions of low and high anisotropy. The range of FA is from 0 (isotropic diffusion) to 1 (highly restricted diffusion) (d) map obtained from the arithmetic average of 26 diffusion-encoding directions, regions of low diffusivity are hyper-intense and regions of high diffusivity are hypo-intense. (e) Mean diffusivity (MD) map, regions of low diffusivity are hypo-intense whereas, regions of high diffusivity are hyper-intense. The range of MD is from 0 to 3×10^−3^ mm^2^/s. Rt: retina; La: lamina; Ax-tk: axonal tracts; MB: mushroom body; Me: medulla; AL: antennal lobe; P: peduncles; Lox: lobula complex.

Using the color encoded projections of synaptic neuropils, it is possible to estimate the micro-diffusion environment (Figure 3). The axonal fibres connecting the synaptic centers of the optic lobes and neuronal projections towards the brain can be seen from Figure 3 (a). Regions that show high contrast in Figure 3 insets 1 – 7 are Rt, MB, Lox, Me, La, CX, AL and Tritocerebral lobe. Similarly, insets 8 -11 show good classification of midbrain neuropil (MN), MB, AL, La, Me and Lox. The central complex (CX), which has high synaptic density appears hypo-intense in the averaged DWI map, whereas, T2-W contrast in the axial plane of Figure 1 shows clear delineation of CX and hypo-intense peduncles (P). The insets in Figure 3 (a) show the 3-dimensional representation of tracts in MB and CX using manually delineated region-specific (ROI-based) fibre tractography. Hence, use of dMRI at micro-scale with multi-contrast maps can be used to identify synapse-rich neuropil sub-structures, cell bodies and axonal tracts.

**Figure 3:**
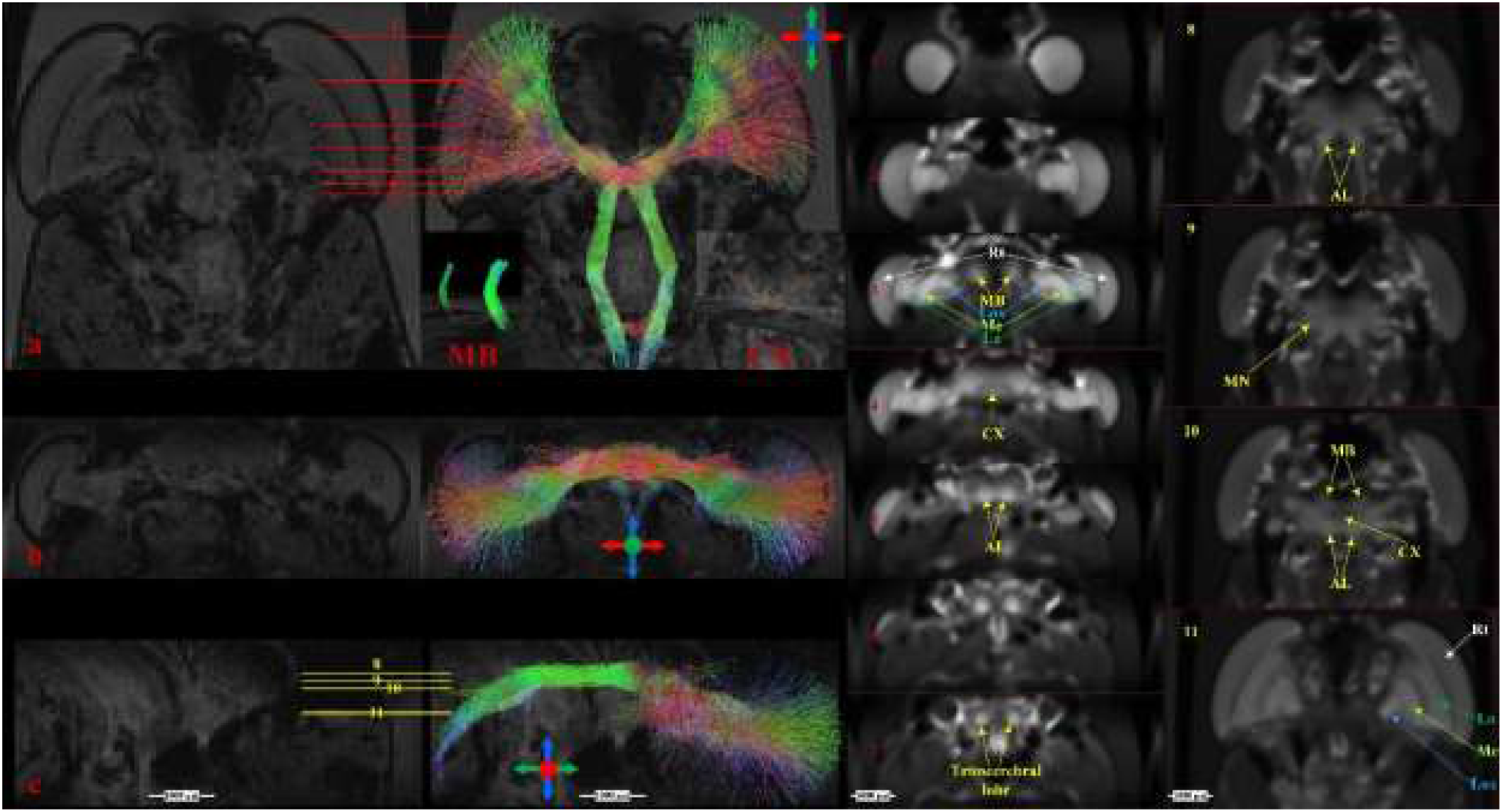
Projections of synaptic neuropils in terms of directionally encoded tracts from sample no. 1 (a) Coronal slice, anterior view, insets 1 – 7 show locations of various anatomical structures in axial plane using the map obtained from the arithmetic average of 26 diffusion-encoding directions. (b) Projection of synaptic pathways in axial plane. (c) Projection of synaptic pathways in Sagittal plane, insets 8 – 11 show locations of various anatomical regions in coronal plane using averaged DWIs. MB: mushroom body CX: central complex; MN: midbrain neuropil

Rotationally invariant measures (FA, MD, MK and KA) provide quantitative information on various micro-structures as can be visualized in an illustrative image from one animal in the coronal (Figure 4) and axial plane (Figure 5). These metrics were extracted for each animal and Rt, La and axonal tracts showed the highest FA values, whereas, CX showed lowest FA (Table 1). Using MK and KA, the complex organization of the underlying microstructure can be quantified. For example, Lox exhibited highest kurtosis values due to its complex micro-structural (diffusion) environment mapped at 78 μm^3^ scale (Table 1), whereas, CX, despite having high synaptic density showed lower kurtosis values as well as low fibre density (Figure 3a-inset), which indicates that much higher resolution is required to better classify high synaptic regions.

**Figure 4:**
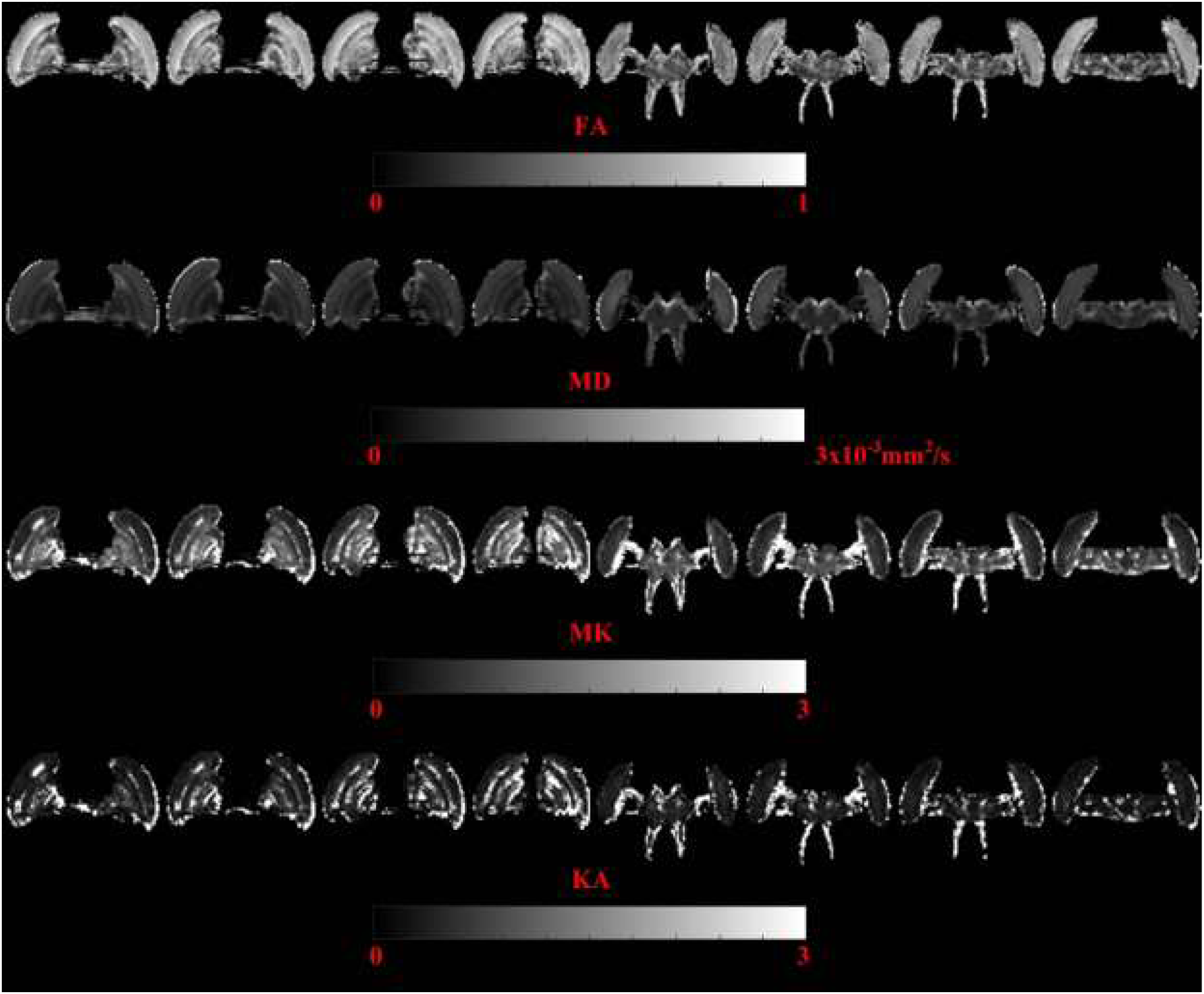
Kurtosis tensor derived directionally invariant indices – coronal plane (sample no. 1). The indices were only calculated in the Regions of interest (ROIs).

**Figure 5:**
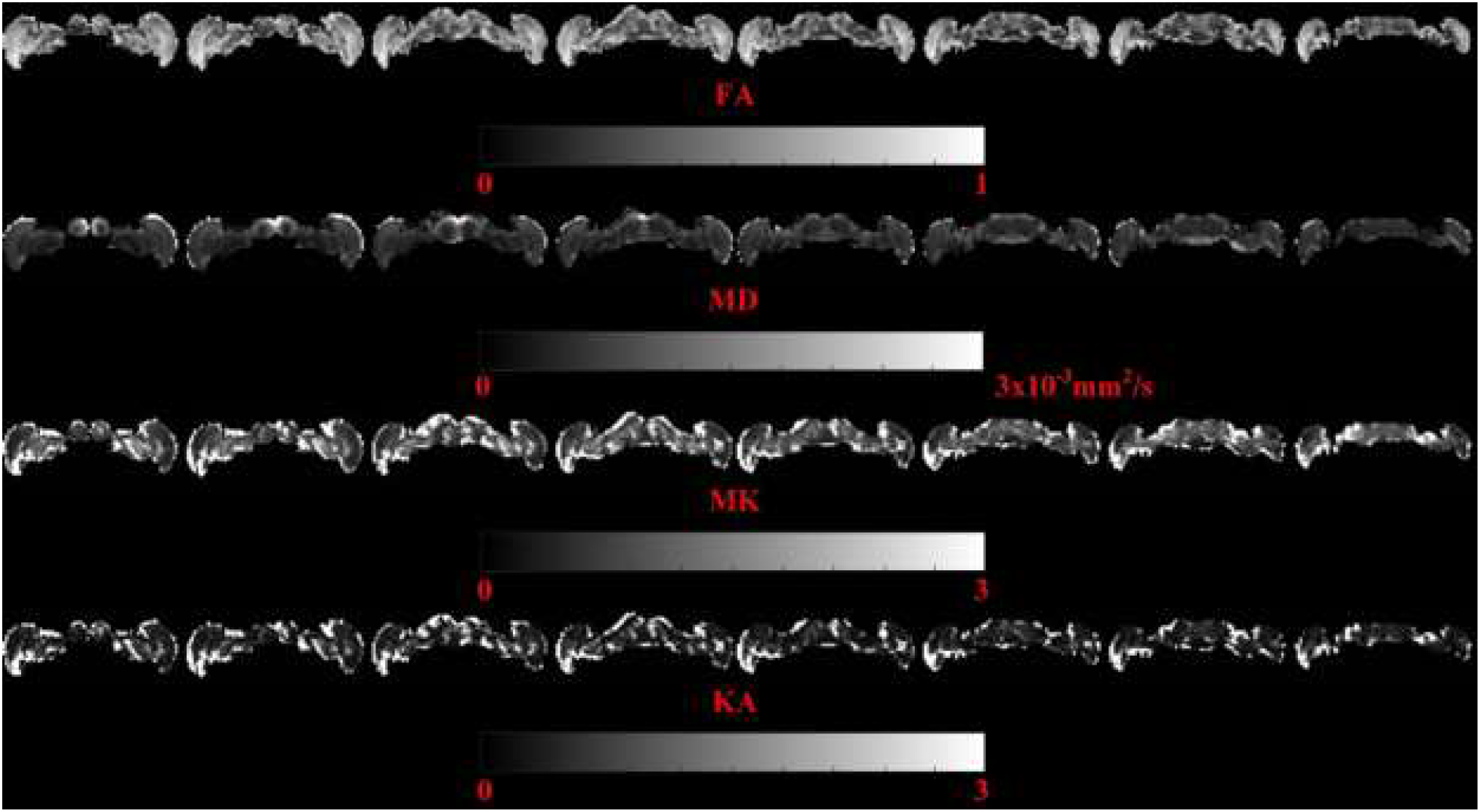
Kurtosis tensor derived directionally invariant indices – axial plane (sample no. 1). The indices were only calculated in the Regions of interest (ROIs).

**Table 1:** median values of Kurtosis tensor derived directionally invariant indices from selected ROIs.

| Locust 1 |  |  |  |  |  |  |  |  |  |
| --- | --- | --- | --- | --- | --- | --- | --- | --- | --- |
|  | AL | CX | Rt | Me | MB | La | MN | Lox | Ax tk |
| **FA** | 0.36 | 0.23 | 0.67 | 0.49 | 0.36 | 0.56 | 0.32 | 0.44 | 0.62 |
| **MD**<br>$\text{m}^2.\text{s}^{-1}$ | $0.46 \times 10^{-03}$ | $0.76 \times 10^{-03}$ | $0.64 \times 10^{-03}$ | $0.35 \times 10^{-03}$ | $0.47 \times 10^{-03}$ | $0.52 \times 10^{-03}$ | $0.47 \times 10^{-03}$ | $0.37 \times 10^{-03}$ | $0.46 \times 10^{-03}$ |
| **MK**<br>$\text{m}^2.\text{s}^{-1}$ | 1.88 | 0.73 | 1.04 | 2.64 | 2.39 | 1.60 | 1.69 | 2.65 | 2.46 |
| **KA** | 0.95 | 0.32 | 0.89 | 1.64 | 1.57 | 0.81 | 0.71 | 1.07 | 1.86 |
| Locust 2 |  |  |  |  |  |  |  |  |  |
| **FA** | 0.50 | 0.31 | 0.68 | 0.49 | 0.50 | 0.67 | 0.51 | 0.62 | 0.67 |
| **MD**<br>$\text{m}^2.\text{s}^{-1}$ | $0.43 \times 10^{-03}$ | $0.62 \times 10^{-03}$ | $0.65 \times 10^{-03}$ | $0.37 \times 10^{-03}$ | $0.36 \times 10^{-03}$ | $0.45 \times 10^{-03}$ | $0.48 \times 10^{-03}$ | $0.26 \times 10^{-03}$ | $0.52 \times 10^{-03}$ |
| **MK**<br>$\text{m}^2.\text{s}^{-1}$ | 2.23 | 0.75 | 0.90 | 2.51 | 3.52 | 2.03 | 2.02 | 4.60 | 2.48 |
| **KA** | 0.92 | 0.39 | 0.59 | 1.11 | 2.08 | 1.19 | 0.93 | 3.83 | 1.60 |
| Locust 3 |  |  |  |  |  |  |  |  |  |
| **FA** | 0.39 | 0.28 | 0.65 | 0.52 | 0.24 | 0.71 | 0.28 | 0.61 | 0.69 |
| **MD**<br>$\text{m}^2.\text{s}^{-1}$ | $0.59 \times 10^{-03}$ | $0.74 \times 10^{-03}$ | $0.57 \times 10^{-03}$ | $0.31 \times 10^{-03}$ | $0.53 \times 10^{-03}$ | $0.41 \times 10^{-03}$ | $0.51 \times 10^{-03}$ | $0.28 \times 10^{-3}$ | $0.42 \times 10^{-03}$ |
| **MK**<br>$\text{m}^2.\text{s}^{-1}$ | 2.24 | 0.85 | 0.89 | 2.04 | 1.94 | 1.84 | 1.77 | 4.91 | 2.65 |
| **KA** | 0.48 | 0.36 | 0.65 | 1.04 | 0.41 | 1.55 | 0.45 | 4.54 | 2.22 |

Interestingly the quantitative metrics for each region were within a similar range for each animal. In future studies for these values to have utility as baseline dMRI metrics there would need to be an increase in sample size, but the initial results are supportive that dMRI metrics could be used to examine structural differences in relation to differing physiology.

## Discussion and Conclusion

There is increasing evidence to support the early observations by Ramon y Cajal which suggested that there were common design principles behind the formation of neuronal networks to subserve complex brain function across phyla ^1,3^. Therefore, the insect brain offers a simpler and more accessible nervous system that can add to understanding of the general principles behind how neuronal networks result in adaptable behavior ^7^. Further the use of insect brains as a model have been proposed to enable better understanding from the molecular level for how ageing ^45^, disease ^46^ or environmental change ^47^ impacts on both neuronal networks and resulting behavioural output. In order to address this, suitable brain imaging techniques are required that will enable the linkage between the underlying anatomy of the neuronal network with functional output, which can be measured via established electrophysiological and behavioural techniques^6^.

This study demonstrates a novel methodology for imaging insect brains. It is demonstrated how dMRI can provide both a qualitative and quantitative insight into the microstructural environment of the locust’s optic lobes and central brain as viewed in an intact head. Regions that have been previously identified by microscopy techniques could be identified in this study ^43^. Previously a number of studies have tried to image the internal organs of various small animals, including insects, but with limited success ^48–50^. These studies mostly employed anatomical (qualitative) scans to observe features such as exoskeleton, guts, ovaries and muscles. Manganese enhanced magnetic resonance imaging (MEMRI); where intracellular accumulation of manganese is used to infer neuronal activity; has been applied to the locust to monitor neuronal activity concurrent with locomotion ^51^. MEMRI can also be applied to trace identifiable neuronal circuits, and this has been successfully applied to *Aplysia* ^52^ but not yet to insects. To the best of our knowledge, there is only one study to date which tried to classify internal structures of insects, (*Drosophila melanogaster*) using FLASH sequence and DWI based contrast ^53^. However, the study only applied diffusion sensitizing gradient along the slice selection gradient. Due to a single diffusion encoding direction and lack of quantitative assessment (apparent diffusion coefficient along slice select direction ADC_Z_), it is difficult in that study to comprehend a more comprehensive diffusion profile of the underlying microstructures. This may be the reason that the authors of that study could not classify axonal tracts and cell bodies. In this study we applied diffusion sensitizing gradients in 26 diffusion encoding directions and used 4 b-values (3 shells). To classify various regions, we used number of contrasts and by employing a Kurtosis tensor we were able to classify the diffusion patterns in neuropils (small axons, dendrites and synaptic terminals) as well as in cell bodies, which are usually difficult to assess using DWI or simple diffusion models.

The application of the dMRI to visualize the insect brain and optic lobes is particularly beneficial for understanding the formation of neuronal circuits and the potential impact of physiological or environmental change on these circuits. In the insect, the visual system is proportionally by far the largest region as may be expected by the necessity of vision for the animal and the demanding computation it requires. There has been extensive prior characterization of the visual system of insects, particularly in flies but also in locusts and bees from the first observation of the similarity in the organization for insect and mammalian visual systems ^44^. The insect visual system is known to be energetically demanding on the animal, with the circuitry highly evolved to efficiently and rapidly transmit information to the central brain^54^. Interestingly there is evidence that there is experience dependent plasticity in the fly optic lobe ^55^. Although the visual system in the insect is already well characterized through microscopy and also electrophysiology, the application of dMRI has the potential to detail dynamic changes that occur in visual system wiring that is necessitated by changing demands on the animal.

The novel application of dMRI and its signal representation schemes based methodology for insect brain imaging is preferable over other possible imaging techniques for many reasons. Non-invasive imaging techniques such as MRI or CT do not interfere with the organization of underlying microstructure, as opposed to histology and microscopic based imaging techniques ^53^. Therefore such techniques are better suited for biological imaging. Additionally, despite the development of processing pipelines for the reconstruction of electron microscopy data ^12^ the technique is time consuming, with this limitation increasing with the size of the neuronal network ^7^. Further, unlike microscopy, the use of dMRI opens up new possible research avenues as the technique could be applied to characterize neuronal connectivity over an animal’s lifespan.

An important addition dMRI provides to the imaging of the insect brain is the availability of dMRI derived quantitative indices (Figures 2 and 4) as well as fibre tractography (Figure 3). DTI metrics including FA have already demonstrated value in human studies of patients with brain trauma or neurodegeneration ^16^. dMRI derived metrics have the potential to quantify changes in the organization of the neuronal microstructure that occurs due to environmental stresses or aging. DTI and DKI are advantageous techniques for characterizing axonal changes as there may not be overt axonal damage even in the presence of significant functional damage ^22^. Therefore better insights into physiological changes in the brain are likely to be gained from this method than through the use of a microscopy technique, even though microscopy has better spatial resolution. With the availability of dMRI in an insect model system there is the potential for a methodology that could be used to study axonal damage from the molecular ^46^ through to the systems level. Recent studies have developed a *Drosophila* model of mild traumatic brain injury (TBI) ^56,57^. Insects, including locusts are good models of TBI as the cellular pathways are highly conserved, the injury can be readily induced and alterations in behaviour easily monitored over the animal’s whole lifespan. After the initial injury, secondary alterations are known to occur in neuronal networks including; membrane depolarization; alterations in neurotransmitter release and activation of voltage dependent ion channels. These changes are followed by activation of conserved inflammatory pathways. Subsequently damaged intracellular material is removed by conserved lysosomal-dependent clearance pathways ^23,57^. Locusts could prove to be particularly good models due to their relatively large brain for insects and the extensive electrophysiological characterization of the neuronal circuits. The advent of gene editing tools such as CRISPR/Cas9 will further increase the utility of the locust to provide mechanistic understanding ^58^

Further applications of dMRI in the insect would be possible if the technique were developed to a *Drosophila* model ^59^. Quantification of alterations in axonal microstructure are important for understanding the mechanisms for the widespread degeneration of axons that is characteristic of many neurodegenerative diseases for which there are existing *Drosophila* models ^22^.

The benefits of dMRI being applied to an insect model may be twofold. Whilst the widely used dMRI derived metrics like FA, KA, MK and MD have been found to have prognostic value in human patients the exact alterations that they are capturing are not completely understood^60,61^. It may be that these metrics are particularly sensitive to a given cellular change, for instance changes in glia. Although there are morphological differences, the insect brain contains glia with analogous functional types to those observed in mammalian brains ^62–64^. Whilst alterations in glia are the most frequently proposed reason for changes in dMRI metrics, axon diameter, packing density and membrane permeability should also be considered ^65,66^. It could be easier in an insect model to determine the mechanistic basis for dMRI derived metrics that may provide valuable understanding when dMRI is applied in other animals, including humans.

There remain some limitations in the dMRI technique. While the DKI is an improvement on the DTI in terms of inferring the underlying biological structure, there is still the possibility for error and tracts can be erroneously inferred or missed. The use of microscopy data in combination with MRI may help to reduce errors in tractography. Technical improvements can reduce the chance of errors in the extraction of tracts, for instance higher dimensionality of the diffusion model or higher field MR. However, although these problems have been inherent in dMRI even as applied to mammalian brains, there is proven value in the utility of dMRI metrics in predicting patient outcomes.

The linkage of metrics from dMRI and DKI with existing well established techniques to characterize functional output of the network provides a powerful methodology to unravel how neuronal structure impacts on behavioural output. This methodology will enable a different and yet complementary approach to understanding the dynamics of neural circuits in development, ageing and stress. Importantly the technique provides the possibility of quantitative as well as qualitative imaging of neuronal structure. The locust is potentially good insect model to develop these studies further as they enable questions to be addressed from a cellular to systems level.

## Data Availability

The raw dMRI volume data used in the study are available from the corresponding author on reasonable request

## Statement of Contributions

S.S.S. and A.G.W. wrote the main manuscript text. S.S.S. performed the data processing and prepared figures 1-4. S.S.S. and CK performed the scans. MB provided critical feedback and guidance on interpretation of results. All authors reviewed the manuscript.

